# Disentangling the Relationship Between Mind Wandering and Symptom Dimensions in a Non-Clinical Sample: The Key Link with ADHD

**DOI:** 10.64898/2026.03.16.712037

**Authors:** Miha Likar, Bianka Brezóczki, Teodóra Vékony, Peter Simor, Dezso Nemeth

## Abstract

**Introduction:** Mind wandering has been linked to a wide range of psychiatric conditions, yet most studies have examined these associations in isolation. Given the substantial comorbidity across the psychopathological spectrum, it remains unclear whether specific symptom dimensions uniquely and independently predict mind wandering once their overlap is accounted for.

**Methods:** To address this, we adopted a dimensional approach in a non-clinical sample (*N* = 384), simultaneously modeling seven symptom dimensions: ADHD, depression, obsessive-compulsive tendencies, schizotypy, autistic traits, hypomania, and eating disorder symptoms.

**Results:** At the bivariate level, mind wandering correlated positively with all symptom dimensions. However, when the substantial shared variance across dimensions was accounted for in both frequentist and Bayesian multivariate regression models, only ADHD symptoms consistently emerged as a unique predictor (β = 0.52; BF₁₀ > 1000). Hypomania symptoms showed a weak, inconsistent positive association across analyses. All remaining predictors yielded negligible unique contributions, with Bayes factors supporting the null hypothesis.

**Conclusion:** These findings suggest that once overlap between symptom dimensions is accounted for, ADHD symptoms remain the only consistent predictor of trait-level mind wandering. We speculate that previously reported associations between mind wandering and other psychopathological symptom dimensions may largely reflect variance shared across the symptom battery rather than dimension-specific mechanisms.

## Introduction

Mind wandering is commonly defined as the shift of attention away from the external environment and ongoing task toward internally generated thoughts [1]. This pervasive cognitive phenomenon [2] is associated with poorer performance on attentionally demanding tasks [3], lower educational attainment [4], and reduced emotional well-being [5], yet paradoxically with enhanced implicit and statistical learning [6, 7]. Mechanistic accounts converge on the view that mind wandering reflects limitations in executive and attentional control, whether framed as a failure to maintain task-relevant focus [8] or as a reallocation of domain-general attentional resources toward internally generated trains of thought [9]. Consistent with these frameworks, higher trait-level mind wandering is associated with elevated levels of attentional and affective symptomatology across multiple dimensions, including ADHD traits [10, 11], depressive symptoms [12], obsessive–compulsive tendencies [13], schizotypal traits [14], and anxiety symptoms [15]. However, most studies have examined these relationships in isolation, with relatively few adopting a dimensional approach that simultaneously models multiple symptom dimensions while accounting for their substantial comorbidity [16–21]. The primary aim of the present study is to examine whether individual psychopathology-related symptom dimensions are associated with mind wandering when their overlap is accounted for. Here, we treat mind wandering not merely as a momentary attentional fluctuation but as a stable cognitive phenotype that varies across individuals and relates to broader neurocognitive traits associated with psychopathology spectra.

The question of whether mind wandering is uniquely linked to specific psychopathology dimensions is particularly pressing because psychopathological symptoms rarely occur in isolation. Substantial comorbidity observed across the psychopathological spectrum [22, 23] has motivated dimensional and transdiagnostic frameworks such as the Hierarchical Taxonomy of Psychopathology [HiTOP; 24, 25] and the NIMH Research Domain Criteria [RDoC; 26, see also 27]. Although a broad range of symptom dimensions has been linked to impairments in executive and attentional control [28–34], accumulating evidence suggests that such executive dysfunctions may constitute a transdiagnostic vulnerability rather than pathology-specific deficits [35–37]. Consequently, given the extensive overlap among symptom dimensions, it remains unclear whether associations between mind wandering propensity and psychopathology-related measures reflect mechanisms specific to particular symptom dimensions or are instead driven by variance shared across dimensions more broadly.

The present study adopts a dimensional approach in a non-clinical sample to examine the relationship between a range of psychopathology-related symptom dimensions and mind wandering. Rather than contrasting diagnostic groups, we examined continuous variation across multiple symptom dimensions, including ADHD symptoms, autism-spectrum traits, depressive symptoms, obsessive–compulsive symptoms, schizotypal traits, hypomanic tendencies, and eating-disorder–related symptoms. We first quantified bivariate associations between mind wandering and each symptom dimension, then assessed their unique contributions in multiple regression models accounting for their overlap. By situating mind wandering within this dimensional model of psychopathology, we explore whether heightened trait-level mind wandering propensity is specific to particular symptom dimensions or broadly shared across psychopathology.

## METHODS

### Participants and procedure

A total of 472 participants were initially recruited through an online experiment in exchange for course credit at Eötvös Loránd University, Hungary. The online experiment was conducted using both the Gorilla Experiment Builder (https://www.gorilla.sc) [38] and Qualtrics platforms. Data collection was conducted in two rounds as part of a larger research project that included additional measures not relevant to the present study. The MWQ was administered via the Gorilla Experiment Builder [38] in both rounds. In the first round, all other questionnaires were also administered via Gorilla, whereas in the second round, the remaining questionnaires were administered via Qualtrics. Thus, although the MWQ response format differed between the two rounds, the platform used for its administration remained the same.

Following quality control procedures, 22 participants were excluded because they had missing complete questionnaire scores for one or more variables of interest, rendering data imputation unfeasible. Eight participants were removed due to incorrect age data, which was a relevant covariate in our analysis. For one participant, a duplicate observation was identified and removed, with the first recorded response retained. Additionally, 57 participants were excluded due to a self-reported mental health diagnosis and/or active use of psychotropic medication. Characteristics regarding the reported diagnoses and medication status of these 57 participants are reported in Supplementary materials S5, Table S6. The final sample used in the analyses consisted of 384 participants (age: *M* = 22.1, *SD* = 4.0, range = 18 - 52; gender: F = 302, M = 80, Other = 2) with complete data across all variables. It should be noted that the absence of a self-reported diagnosis or medication use in this final sample does not imply the absence of psychopathology-related symptoms. Instead, the “non-clinical” label refers specifically to the lack of a self-reported formal diagnosis or associated treatment. Participants provided informed consent, and the studies received approval from the Research Ethics Committee of Eötvös Loránd University, Budapest, Hungary (2024/214), in accordance with the principles of the Declaration of Helsinki.

### Instruments

The **Mind-Wandering Questionnaire** [MWQ; 39] is a brief self-report questionnaire assessing an individual’s trait-level propensity to engage in mind wandering. The Hungarian adaptation of the MWQ contains 5 items [40]. In its original form, MWQ items are rated on a 6-point Likert scale (range 5-30). However, in the present study, part of the sample completed the questionnaire using an 8-point Likert scale (range 0-35) due to a change in the response format during data collection.

The **Autism–Spectrum Quotient** [ASQ; 41] is a 50-item self-report scale that measures traits associated with the autism spectrum in the adult population. The scale assesses five symptom clusters of the autism spectrum where items are rated on a 4-point Likert scale (range 0-50).

The **Adult ADHD Self-Report Scale** [ASRS; 42], an 18-item self-report scale, was employed to assess ADHD symptom severity. The ASRS provides a total score (range 0-72) and two subscale scores with 9 items measuring *Inattention* and 9 measuring *Hyperactivity/Impulsivity*. Responses are indicated on a 5-point Likert scale.

The **Beck Depression Inventory** [BDI; 43] is a widely used 21-item self-report measure of depressive symptom severity. We used the 9-item Hungarian short form [44, 45], which assesses core depressive symptoms using items derived from the original scale. Items were rated on a 4-point response scale ranging from 1 (“not typical”) to 4 (“very typical”), yielding a total score between 9 and 36. Psychometric evaluation of the Hungarian short form supported a unidimensional factor structure and demonstrated criterion validity against depression diagnoses established using a structured diagnostic interview [45]. Although originally validated in an adult primary-care sample, the 9-item version has also been applied in population-based, non-clinical Hungarian samples [46].

The **Eating Attitude Test** [EAT; 47] is a self-report scale assessing the symptoms and concerns related to eating disorders. A 26-item shortened version was applied, assessing dieting behaviors, bulimia, and food preoccupation, and oral control on a 6-point Likert scale (range 0-78).

The **Hypomania Checklist** [HCL-32; 48], a self-report questionnaire consisting of 32 items, was used to assess hypomanic symptoms. Questions concern two factors, one evaluating active/elated mood and the other rating irritable/risk-taking tendencies on a binary (Yes/No) scale (range 0-32).

A brief version of the **Multidimensional Schizotypy Scale** [MSS-B; 49] was used to assess schizotypal traits. Using a binary (Yes/No) scale on 38 items, this questionnaire measures the *positive*, *negative,* and *disorganised* dimensions of schizotypal personality traits in non-clinical populations (range 0-38).

**Obsessive–Compulsive Inventory-Revised** [OCI-R; 50] was employed to assess OCD symptoms. The scale consists of 18 items, measuring symptoms across 6 domains – washing, obsessing, hoarding, ordering, checking, and neutralizing. Items are rated on a 5-point scale (range 0-72).

For all scales, higher scores indicate greater symptom severity. All questionnaires demonstrated good to excellent internal consistency (see Table 1 for scale-specific values).

**Table 1:** Descriptive statistics and internal consistency for all study variables.

| Questionnaire | Mean (SD) | Skewness | Kurtosis | Cronbach's $\alpha$ |
| --- | --- | --- | --- | --- |
| MWQ (mind wandering) (Round 1) | 19.28 (7.28) | −0.52 | 0.06 | 0.88 |
| MWQ (mind wandering) (Round 2) | 19.91 (5.20) | −0.11 | −0.56 | 0.82 |
| ASQ (autism spectrum) | 19.02 (5.74) | 0.32 | −0.27 | 0.79 |
| ASRS (ADHD) | 50.23 (9.54) | 0.12 | 0.10 | 0.84 |
| BDI shortened (depression) | 14.52 (4.91) | 0.74 | −0.46 | 0.87 |
| EAT (eating attitude) | 11.88 (11.61) | 1.62 | 2.59 | 0.90 |
| HCL-32 (hypomania) | 17.98 (6.19) | −0.67 | 0.56 | 0.87 |
| MSS-B (schizotypy) | 8.47 (6.74) | 0.95 | 0.28 | 0.89 |
| OCI-R (OCD) | 20.27 (12.88) | 0.38 | −0.74 | 0.91 |

*Note.* Mean scores, standard deviations, skewness, kurtosis, and Cronbach's $\alpha$ coefficients are presented for each questionnaire. MWQ statistics are reported separately by data collection round due to different response scales (Round 1: N = 143; Round 2: N = 241). ASQ = Autism Spectrum Quotient; ASRS = Adult ADHD Self-Report Scale; BDI = Beck Depression Inventory; EAT = Eating Attitude Test; HCL-32 = Hypomania Checklist; MSS-B = Multidimensional Schizotypy Scale-Brief; MWQ = Mind-Wandering Questionnaire; OCI-R = Obsessive-Compulsive Inventory-Revised.

### Data analysis

All data analyses and visualisations were conducted in R [version 4.4.2;, 51] using packages *tidyverse* [52] and *sjPlot* [53] for preprocessing and visualisation, respectively. All variables were standardized prior to analysis. To account for differences in response scale format, MWQ scores were standardized separately within each data collection round (Round 1: N = 143; Round 2: N = 241), such that scores from both rounds were expressed on a common metric (M = 0, SD = 1) before being combined. Importantly, the two rounds did not differ in any other procedural respect, and the standardization approach ensures that scale format differences do not introduce systematic bias into the analyses. Statistical significance was evaluated at α = .05.

To explore zero-order correlations between all cross-sectional questionnaires, bivariate Spearman’s rank correlations were computed using all available observations (N = 384 for each respective correlation). The false discovery rate (FDR) procedure was used for p-value adjustment to account for multiple comparisons. Given the exploratory nature of the correlation analysis, this approach was chosen over more conservative approaches to control the expected proportion of false positives while maintaining statistical power. Correlation coefficients (ρ) and FDR-adjusted p-values are reported. Correlation analysis was performed using *psych* [54].

To investigate the unique contribution of respective psychopathology-related questionnaires to mind wandering scores while accounting for their shared variance, multiple linear regression analyses were conducted. Mind wandering was entered as the dependent variable, psychopathology-related questionnaire scores were included as predictors, with age and gender entered as covariates. According to formal testing and visualisations conducted with the *performance* package [55], the model did not violate assumptions of normality, homoscedasticity, or outlier influence. Multicollinearity was not a concern, with all variance inflation factors (VIFs) below 1.93. Standardized regression coefficients (β), standard errors, confidence intervals, semi-partial r² calculated using *effectsize* [56], and model fit indices are reported.

To complement the frequentist regression analysis with a probabilistic estimation of parameter uncertainty and to quantify evidence for model parameters, Bayesian linear regression was additionally conducted using the *brms* package [57]. Weakly informative priors were used (Normal(0, 1) for regression coefficients and intercept), which provide slight regularization without strongly constraining the estimates. To assess the robustness of the model, sensitivity analysis was performed with moderately conservative (Normal(0, 0.5)) and strongly regularizing (Normal(0, 0.3)) priors for regression coefficients (*see **Supplementary Materials S4***). Intercept priors (Normal(0, 1)) were identical across all specifications. Standardized regression coefficients (β) as posterior means, standard errors of the posterior distribution (Est. Error), 95% credible intervals (CrIs), and Bayes factors (BF10) indicating the strength of evidence for each predictor’s effect are reported [58]. Bayes factors were computed using *bayestestR* [59]. Bayesian R² is reported as a measure of overall model fit.

Additional sensitivity analyses were conducted to further probe the robustness of associations observed in the primary regression model. These are reported alongside the relevant results, with full model outputs provided in Supplementary Materials S1, S2 and S3. To assess the robustness of the main findings, all main analyses were additionally conducted on an extended dataset that retained participants with a self-reported mental health diagnosis and/or active use of psychotropic medication (*N* = 441). Results of these analyses are reported in Supplementary Materials S5.

## RESULTS

### Descriptive statistics

Descriptive statistics and Cronbach’s α values for all variables of interest are reported in Table 1. For MWQ, descriptive statistics are reported separately for the two subsamples corresponding to different data collection rounds. Inspection of skewness and kurtosis values indicated approximately normal distributions for most scales. EAT scores showed moderate positive skew and leptokurtosis. All questionnaires demonstrated good to excellent internal consistency, with Cronbach’s α ranging from 0.79 (ASQ) to 0.91 (OCI-R).

### Mind wandering is positively associated with diverse psychopathology-related symptoms

Full results of Spearman’s rank bivariate correlations are reported in the correlation matrix in Figure 1. They revealed that mind wandering showed a moderate to strong positive association with ADHD symptoms (ρ = .52, *pFDR* < .001, N = 384), and weak to moderate positive associations with depressive symptoms (ρ = .26, *pFDR* < .001, N = 384), schizotypal traits (ρ = .26, *pFDR* < .001, N = 384), and obsessive-compulsive symptoms (ρ = .26, *pFDR* < .001, N = 384). Weak but significant associations were observed between mind wandering and autistic traits (ρ = .12, *pFDR* < .05, N = 384), hypomania symptoms (ρ = .18, *pFDR* < .001, N = 384), and eating attitudes (ρ = .13, *pFDR* < .05, N = 384). These results indicate that, at the bivariate level, higher trait-level mind wandering propensity is positively associated with a range of psychopathology-related symptoms.

**Figure 1:**
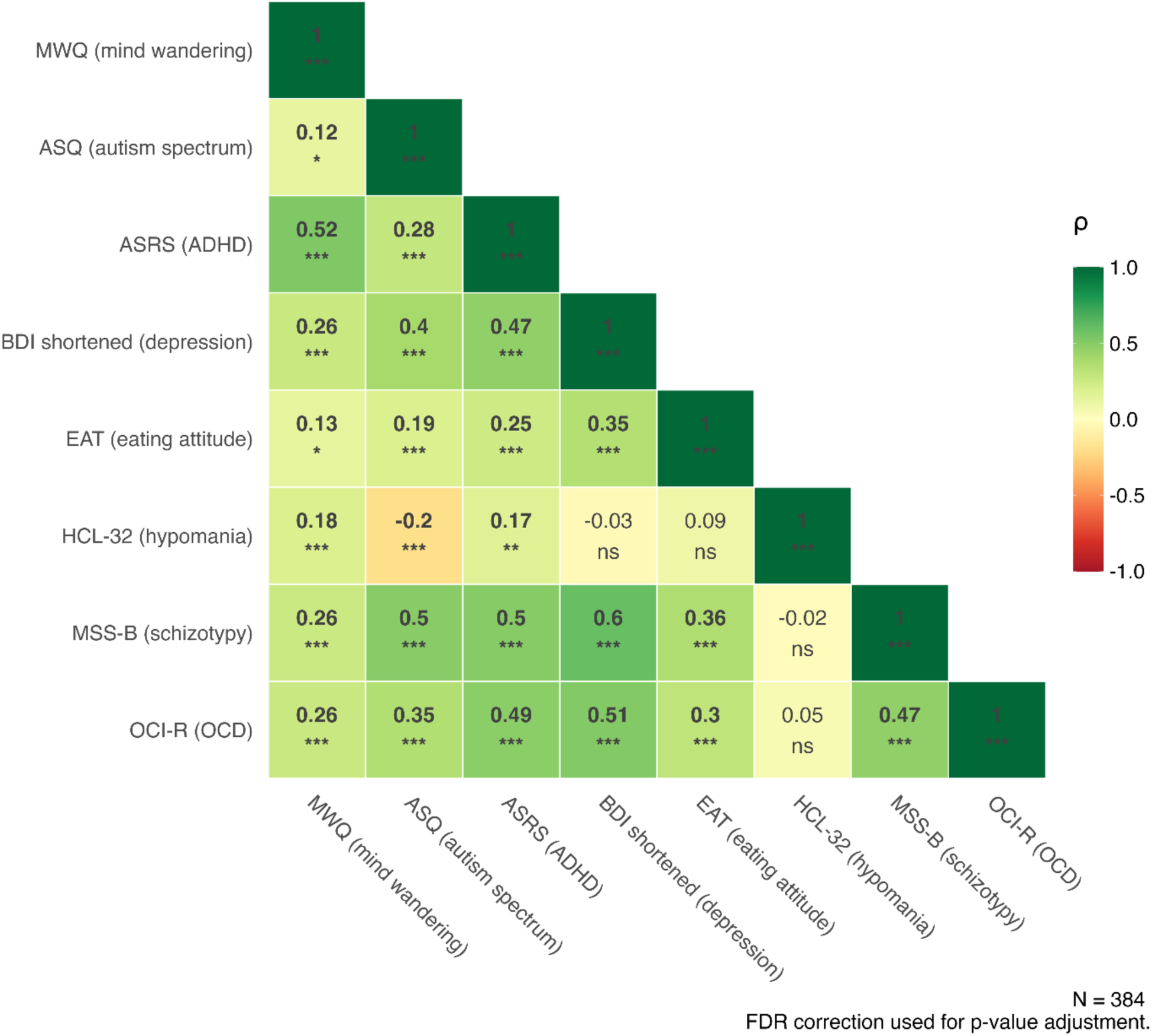
Correlations among mind wandering and psychopathology-related measures. *Note.* ASQ = Autism Spectrum Quotient; ASRS = Adult ADHD Self-Report Scale; BDI = Beck Depression Inventory; EAT = Eating Attitude Test; HCL-32 = Hypomania Checklist; MSS-B = Multidimensional Schizotypy Scale-Brief; MWQ = Mind-Wandering Questionnaire; OCI-R = Obsessive-Compulsive Inventory-Revised. * p < .05; ** p < .01; *** p < .001 (FDR-corrected).

Importantly, significant moderate to strong associations were also observed across the psychopathological symptoms themselves, particularly between ADHD, depressive, schizotypal, obsessive-compulsive, and autistic traits and symptoms. Other associations were weaker or non-significant and are reported in detail in Figure 1. These results reveal a substantial symptom overlap across psychopathology-related measures of interest.

### Mind wandering shows the strongest association with ADHD-like symptoms

The multiple regression model was statistically significant (*F_(10, 373)_ =* 16.84*, p <* .001), explaining approximately 31% of the variance in mind wandering scores (adjusted *R*² = .29). ASRS scores, indicating ADHD symptom severity, emerged as the strongest significant predictor of mind wandering (β = 0.52, *SE =* 0.05*, t =* 9.56*, p* < .001) (Figure 2, panels A and B). HCL-32 scores, indicating hypomania symptoms, emerged as a significant positive predictor (β = 0.10, *SE =* 0.05*, t =* 2.08*, p* = .038). All other predictors, including depressive symptoms, schizotypal traits, obsessive-compulsive symptoms, autistic traits, and eating disorder symptoms, did not emerge as significant predictors after accounting for the remaining symptom dimensions (full results are reported in Table 2). Semi-partial r² indicated that ASRS scores uniquely accounted for 17% of the variance in mind wandering scores. HCL-32 scores, while statistically significant, accounted for less than 1% of the variance in mind wandering scores. All other predictors each accounted for less than 1%. This pattern contrasts with the bivariate correlations reported above, where most symptom dimensions showed significant associations with mind wandering (see Figure 1). In the multiple regression model, these associations were substantially attenuated, with ADHD symptoms remaining the strongest predictor and HCL-32 showing a comparatively weak association.

**Figure 2:**
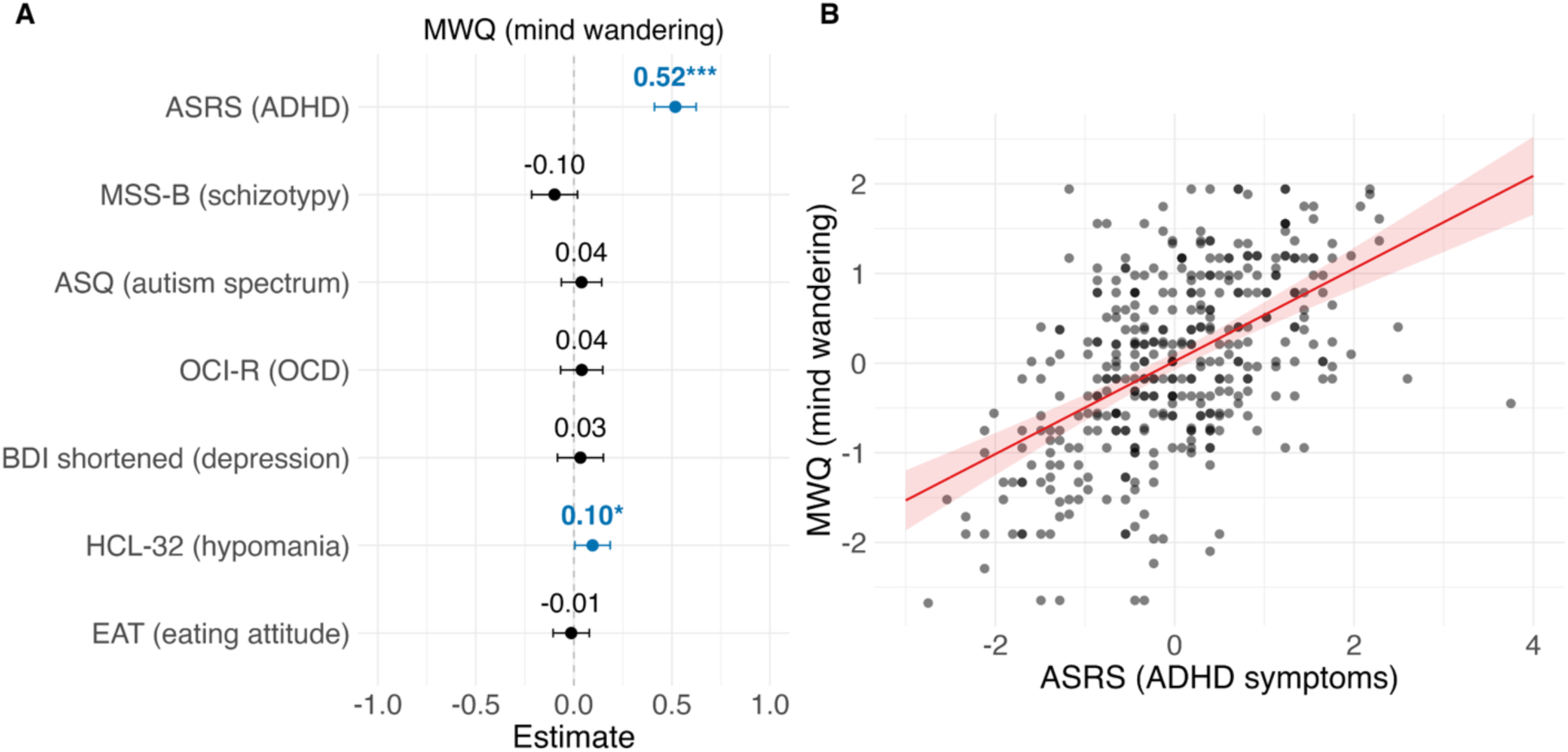
Multiple linear regression predicting mind wandering from psychopathology-related measures. *Note.* **Panel A:** Standardized regression coefficients (β) with 95% confidence intervals for all predictors. The vertical dashed line represents no effect (β = 0). **Panel B:** Model-based predicted mind wandering scores as a function of ADHD (ASRS), estimated from a multiple regression model. Predictions are shown with all other predictors held constant at their mean. Individual data points represent standardized scores, the red line shows the predicted linear relationship, and the shaded ribbon represents the 95% confidence interval around the prediction line. ASRS = Adult ADHD Self-Report Scale; MSS-B = Multidimensional Schizotypy Scale-Brief; ASQ = Autism Spectrum Quotient; OCI-R = Obsessive-Compulsive Inventory-Revised; BDI = Beck Depression Inventory; HCL-32 = Hypomania Checklist; EAT = Eating Attitude Test; MWQ = Mind-Wandering Questionnaire. * p < .05; ** p < .01; *** p < .001

**Table 2:** Multiple linear regression predicting mind wandering from psychopathology-related measures.

|  | **MWQ (mind wandering)** |  |  |  |  |  |  |
| --- | --- | --- | --- | --- | --- | --- | --- |
| *Predictors* | $\beta$ | *SE* | *CI* | *t-value* | *p* | *df* | *sr*<sup>2</sup> |
| (Intercept) | 0.02 | 0.05 | -0.08 – 0.12 | 0.42 | 0.678 | 373.00 | - |
| ASRS (ADHD) | 0.52 | 0.05 | 0.41 – 0.62 | 9.56 | **<0.001** | 373.00 | .17 |
| MSS-B (schizotypy) | -0.10 | 0.06 | -0.22 – 0.02 | -1.66 | 0.098 | 373.00 | <.01 |
| ASQ (autism spectrum) | 0.04 | 0.05 | -0.06 – 0.14 | 0.75 | 0.455 | 373.00 | <.01 |
| OCI-R (OCD) | 0.04 | 0.05 | -0.07 – 0.15 | 0.73 | 0.463 | 373.00 | <.01 |
| BDI shortened (depression) | 0.03 | 0.06 | -0.08 – 0.15 | 0.56 | 0.574 | 373.00 | <.01 |
| HCL-32 (hypomania) | 0.10 | 0.05 | 0.01 – 0.19 | 2.08 | **0.038** | 373.00 | <.01 |
| EAT (eating attitude) | -0.01 | 0.05 | -0.11 – 0.08 | -0.28 | 0.776 | 373.00 | <.01 |
| Age | 0.00 | 0.04 | -0.08 – 0.09 | 0.10 | 0.917 | 373.00 | <.01 |
| Gender [male] | -0.12 | 0.11 | -0.34 – 0.09 | -1.11 | 0.267 | 373.00 | <.01 |
| Gender [other] | 1.03 | 0.60 | -0.15 – 2.20 | 1.71 | 0.088 | 373.00 | <.01 |
| Observations | 384 |  |  |  |  |  |  |
| R<sup>2</sup> / R<sup>2</sup> adjusted | .31 / .29 |  |  |  |  |  |  |
| F<sub>(10, 373)</sub> = 16.84, p < .001 |  |  |  |  |  |  |  |

*Note.* Standardized regression coefficients ( $\beta$ ), standard errors (SE), 95% confidence intervals (CI), t-values, p-values, degrees of freedom, and semi-partial $r^2$ are reported for each predictor. p-values of significant predictors are indicated in bold. ASRS = Adult ADHD Self-Report Scale; MSS-B = Multidimensional Schizotypy Scale-Brief; ASQ = Autism Spectrum Quotient; OCI-R = Obsessive-Compulsive Inventory-Revised; BDI = Beck Depression Inventory; HCL-32 = Hypomania Checklist; EAT = Eating Attitude Test. Gender was dummy-coded with female as the reference category.

To evaluate the robustness of the association between ADHD symptoms and mind wandering, additional supplementary sensitivity analyses were conducted. First, the ASRS Inattention and Hyperactivity/Impulsivity subscales (ρ = .56, p < .001) were entered simultaneously instead of the ASRS total score to examine whether the observed association was specific to the attentional symptom subscale. Both subscales showed significant associations with mind wandering, though the association was notably stronger for the Inattention subscale (β = 0.43, *SE* = 0.06, *t* = 7.51, *p* <.001, *sr²* = .10) than for the Hyperactivity/Impulsivity subscale (β = 0.16, *SE* = 0.06, *t* = 2.87, *p* = .004, *sr²* = .01). Full model output is reported in Supplementary Materials S1.

Second, given the notable semantic overlap in how the ASRS and MWQ assess ADHD symptoms and mind wandering, respectively, we additionally explored the contribution of overlapping items to the observed association between ADHD and mind wandering. Using correlation analysis and content comparison, we identified the three ASRS items (items 7, 8, and 9) with the closest semantic overlap with MWQ content. Averaged across the five MWQ items, these three items showed the highest mean correlations with MWQ items (r = .32, .44, and .31). We examined their contribution to the association in two analyses. In the first, we excluded the identified overlapping items from the ASRS total score. This revealed that the association between ADHD symptoms and mind wandering was attenuated but remained significant (β = 0.42, *SE* = 0.05, *t* = 7.73, *p* < .001, *sr²* = .12), compared to the original association observed with the full ASRS score (β = 0.52). In the second, we entered the overlapping items alongside the remaining ASRS items simultaneously. This revealed that both showed significant independent associations with mind wandering (overlapping items: β = 0.46, *SE* = 0.05, *t* = 8.40, *p* <.001, *sr²* = .12; remaining items: β = 0.20, *SE* = 0.06, *t* = 3.52, *p* < .001, *sr²* = .02), with the overlapping items carrying substantially greater weight per item, given they comprised only three of the eighteen ASRS items. Identified overlapping ASRS items and full model output for both analyses are reported in Supplementary Materials S2.

Lastly, given that the MWQ response format differed across the two data collection rounds, we tested an ASRS × Round interaction to assess whether the ADHD-mind wandering association was consistent across rounds. The interaction was non-significant (β = 0.10, *SE* = 0.09, *t* = 1.10, *p* = .271), and round-specific slopes were comparable (Round 1: β = 0.46, 95% CI [0.30, 0.61]; Round 2: β = 0.56, 95% CI [0.43, 0.69]), indicating that the association between ADHD symptoms and mind wandering did not differ meaningfully across data collection rounds. Full results of this analysis are reported in Supplementary Materials S3.

Results of the regression model on the extended dataset (including diagnosed or medicated individuals) are reported in **Supplementary Materials S5.3.** Although HCL-32 reached statistical significance in this model, its semi-partial r² value was negligible (*sr²* < .01). The overall pattern of results thus remains consistent with the main analysis, with ADHD symptom severity being the strongest significant predictor of mind wandering.

### Bayesian regression

Model convergence was satisfactory (all R^ = 1.00), with effective sample sizes indicating stable posterior estimation. As shown in Table 3, ADHD symptoms showed strong evidence for a positive association with mind wandering (β = 0.52; 95% CrI [0.41, 0.62]). The Bayes factor indicated decisive evidence in favor of the alternative hypothesis (BF₁₀ > 1000). In contrast to the frequentist regression, the Bayesian analysis provided only weak, inconclusive evidence for the HCL-32 (hypomania) association. Although the 95% credible interval excluded zero [0.01, 0.18], the Bayes factor (BF₁₀ = 0.40) indicated evidence favoring the null over the alternative hypothesis. Consistent with the frequentist regression analysis, no other psychopathology-related measures showed evidence for unique associations with mind wandering. For all remaining predictors, 95% credible intervals included zero, and Bayes factors provided evidence in favor of the null hypothesis over the alternative (BF₁₀ < 1), except for the undefined gender category, for which the Bayes factor suggested anecdotal evidence (BF₁₀ = 1.48). This result should be interpreted with extreme caution, as only two individuals belonged to this group. Overall, the model explained a substantial proportion of variance in mind wandering (Bayesian R² = .31, 95% CrI [0.25, 0.37]). Sensitivity analyses with more conservative priors yielded consistent results (*see Supplementary S4*).

**Table 3.**
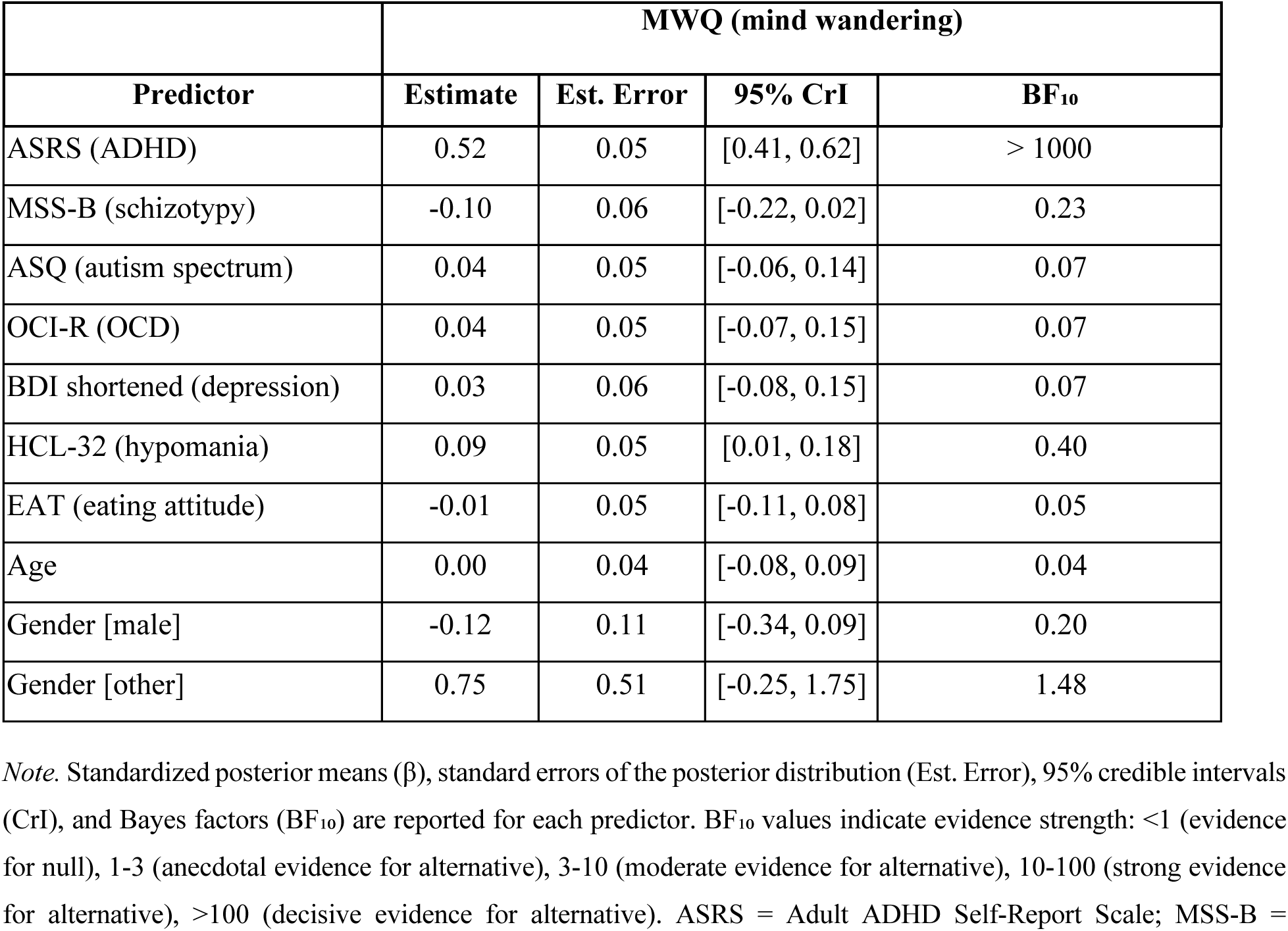
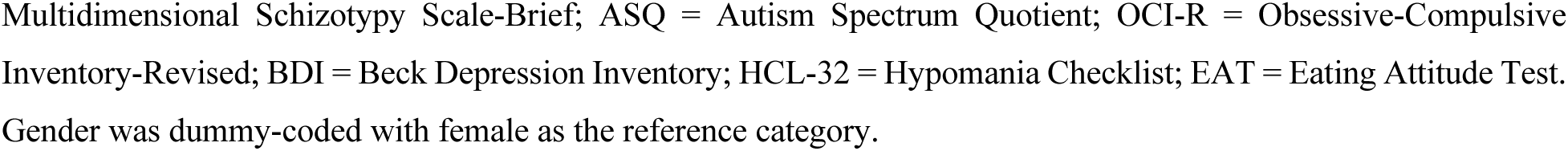
Bayesian multiple linear regression predicting mind wandering from psychopathology-related measures.

|  | **MWQ (mind wandering)** |  |  |  |
| --- | --- | --- | --- | --- |
| **Predictor** | **Estimate** | **Est. Error** | **95% CrI** | **BF<sub>10</sub>** |
| ASRS (ADHD) | 0.52 | 0.05 | [0.41, 0.62] | > 1000 |
| MSS-B (schizotypy) | -0.10 | 0.06 | [-0.22, 0.02] | 0.23 |
| ASQ (autism spectrum) | 0.04 | 0.05 | [-0.06, 0.14] | 0.07 |
| OCI-R (OCD) | 0.04 | 0.05 | [-0.07, 0.15] | 0.07 |
| BDI shortened (depression) | 0.03 | 0.06 | [-0.08, 0.15] | 0.07 |
| HCL-32 (hypomania) | 0.09 | 0.05 | [0.01, 0.18] | 0.40 |
| EAT (eating attitude) | -0.01 | 0.05 | [-0.11, 0.08] | 0.05 |
| Age | 0.00 | 0.04 | [-0.08, 0.09] | 0.04 |
| Gender [male] | -0.12 | 0.11 | [-0.34, 0.09] | 0.20 |
| Gender [other] | 0.75 | 0.51 | [-0.25, 1.75] | 1.48 |

*Note.* Standardized posterior means ( $\beta$ ), standard errors of the posterior distribution (Est. Error), 95% credible intervals (CrI), and Bayes factors ( $BF_{10}$ ) are reported for each predictor. $BF_{10}$ values indicate evidence strength: <1 (evidence for null), 1-3 (anecdotal evidence for alternative), 3-10 (moderate evidence for alternative), 10-100 (strong evidence for alternative), >100 (decisive evidence for alternative). ASRS = Adult ADHD Self-Report Scale; MSS-B =
Multidimensional Schizotypy Scale-Brief; ASQ = Autism Spectrum Quotient; OCI-R = Obsessive-Compulsive Inventory-Revised; BDI = Beck Depression Inventory; HCL-32 = Hypomania Checklist; EAT = Eating Attitude Test. Gender was dummy-coded with female as the reference category.

In the Bayesian analyses of the extended dataset (*see Supplementary materials S5.4*), ADHD symptoms remained the strongest predictor across all prior specifications, with BF₁₀ > 1000 throughout. In contrast, HCL-32 showed prior-dependent evidence ranging from ambiguous to anecdotal-to-moderate (BF₁₀ = 0.93–3.10 across prior specifications), with evidence strength increasing under more regularizing priors. The inclusion of diagnosed and medicated individuals did not alter the primary pattern of results.

## Discussion

The present study investigated the associations between mind wandering and various psychopathological symptoms, adopting a dimensional approach and modeling multiple domains simultaneously in a non-clinical sample. We observed that mind wandering was associated with a broad range of psychopathology-related symptoms at the zero-order level, including ADHD symptoms, autism-spectrum traits, depressive symptoms, OCD symptoms, schizotypal traits, hypomanic tendencies, and eating-disorder–related symptoms. Importantly, significant correlations were observed among the psychopathology-related measures themselves. When this shared variance among psychopathological measures was taken into account in frequentist and Bayesian multiple regression models, only ADHD symptoms remained consistently associated with mind wandering. HCL-32, indicating hypomania symptoms showed an inconsistent pattern of association across analyses. We return to this finding after our discussion of the ADHD-mind wandering association below.

Significant bivariate correlations between the psychopathology-related scales and mind wandering are consistent with prior work linking elevated mind wandering to diverse forms of psychopathology, including ADHD traits [10, 11], depressive symptoms [12], obsessive–compulsive tendencies [13], and schizotypal traits [14]. Interestingly, these associations were substantially attenuated for every questionnaire except the ASRS once all seven dimensions were modeled simultaneously in multiple regression, where shared variance was controlled for. One possible - albeit speculative - account for this attenuation is that it reflects an underlying common factor shared across the whole battery, potentially contributing to the observed bivariate correlations. The idea of common factor would be consistent with dimensional accounts of psychopathology that emphasize a general psychopathology factor (”p-factor”), capturing variance shared across diverse symptom domains [22]. Some accounts link this factor to executive or attentional control liabilities common to multiple symptom dimensions [35, 37], though its interpretation remains actively debated [60, 61]. A shared psychopathology-related factor could therefore provide one explanation for the pattern of attenuated associations observed when the symptom dimensions were modeled simultaneously. Whether such shared variance reflects attentional or executive dysfunction, however, cannot be determined from our findings. The present modeling approach does not allow us to isolate such a common factor, define its nature, or assess its relationship with mind wandering directly. This interpretation should therefore be understood as a theoretical speculation rather than a demonstrated finding.

Our modeling approach should also be considered when interpreting the significant association between ADHD symptoms and mind wandering. ASRS and the other symptom measures are by nature imperfect proxies, and each measure captures a mixture of both a domain-specific and a shared component, exhibiting substantial correlations. Entering them simultaneously in a regression model substantially reduces, but cannot fully eliminate, the variance a given measure shares with the rest of the battery. Correlation between measures did not lead to multicollinearity problems in multiple regression models. The resulting regression coefficients are best interpreted as conditional effects of different measurement scales, indexing each symptom measure’s association with mind wandering while controlling for the others. They should not be directly interpreted as evidence for cleanly separated underlying constructs or mechanistically distinct processes. A significant ASRS coefficient cannot be assumed to reflect exclusively ADHD-specific variance, and at least two alternative explanations should be considered.

First, ADHD symptoms may show genuinely specific associations with mind wandering, over and above what they share with the other symptom dimensions, arguably reflecting specific attentional-regulation difficulties characteristic of ADHD. Second, ADHD symptoms may be a particularly strong indicator of the broader shared psychopathology-related factor discussed earlier. These explanations are not mutually exclusive, and both shared and ADHD-specific components may contribute to the observed association. Future studies employing larger sample sizes and latent-variable approaches could explicitly model this shared structure to better disentangle general and dimension-specific contributions to mind wandering. Inclusion of direct measures of attentional and executive functioning could further test whether objectively assessed attentional-control difficulties may partly account for shared variance.

The dominant association between ADHD symptoms and mind wandering observed here is broadly consistent with theoretical accounts arguing that mind wandering reflects limitations in attentional and executive functioning [8]. Moreover, our findings are also consistent with prior studies adopting multivariate approaches at modeling psychopathological dimensions and mind wandering. For example, Moukhtarian et al. [19] found that when controlling for depression and anxiety, only individuals with ADHD continued to show elevated spontaneous mind wandering relative to controls, while differences for other clinical groups disappeared. Similarly, Gionet et al. [17] observed that inattention symptoms uniquely predicted trait-level mind wandering when modeled simultaneously with cognitive disengagement syndrome and internalizing symptoms. Ogata et al. [20] reported that ADHD independently predicted mind wandering in adolescents even after controlling for autistic traits and depressive symptomatology. Across these studies, a consistent pattern emerges: when symptom dimensions are modeled jointly, ADHD-related symptoms reliably emerge as the dominant predictor of trait-level mind wandering. By including an even broader set of psychopathology-related dimensions, the present study extends these findings and suggests that, among the symptom domains examined here, the association with mind wandering is most strongly concentrated in ADHD symptomatology. Importantly, the core pattern of results remained consistent when analyses were extended to include participants reporting psychiatric diagnoses or psychotropic medication use, with ADHD symptoms retaining their strongest association with mind wandering.

Notably, our results appear to diverge from Figueiredo et al. [16], who found that mind wandering was significantly associated with anxiety but observed no differences between individuals with and without ADHD. This discrepancy may reflect critical methodological differences: Figueiredo et al. examined a smaller clinical sample of diagnosed adolescents with substantial comorbidity, in which anxiety symptoms may be particularly prominent, whereas our study assessed dimensional variation across a broader range of psychopathology-related symptoms in a larger non-clinical sample. More broadly, comparisons between our findings and those from clinical or diagnosed samples [e.g., 16, 19, 20] should be made with caution given this design difference. We view our non-clinical, dimensional approach as a strength, as it may reduce confounds related to diagnostic status, treatment, and clinical severity. Nonetheless, whether the present associations generalize to clinical populations and formal diagnostic groups remains an open question for future research.

Given the significance of ADHD in the present findings, it is worth briefly contextualizing this association within neurobiological accounts linking mind wandering and ADHD. One prominent account [10] proposes that excessive, spontaneous mind wandering in ADHD arises from dysfunctional interaction between two large-scale brain networks. First, the default mode network fails to efficiently deactivate during task performance, and second, the fronto-parietal executive control network shows reduced recruitment during task engagement in ADHD. This account speculates that mind wandering and ADHD-related inattentiveness may share a common underlying mechanism rooted in dysfunctional default mode network activity and connectivity. This raises the possibility that the two constructs partly index the same thing [10]. A second, more recent line of evidence points to local sleep activity - brief, sleep-like slow waves that occur in specific regions of the cortex during wakefulness and have been linked to both mind wandering and mind blanking [62–64]. This activity occurs at a higher density in adults with ADHD than in neurotypical individuals and could help explain their heightened mind wandering [65]. Genetic evidence offers a different, more distal kind of support for treating ADHD as distinct among the symptom dimensions examined here. Large-scale genetic analyses position ADHD separately from many other psychiatric disorders, which cluster more within compulsive or internalizing dimensions [66]. This distinct genetic positioning is broadly consistent with ADHD’s dominant association with mind wandering observed in the present study. It raises the possibility that the association reflects neurodevelopmental attentional deficits distinct from mechanisms underlying compulsive behavior or emotional dysregulation, which are more characteristic of other symptom dimensions. However, as none of these neurobiological accounts were directly tested in the present study, they are offered here as broader context for the present results, rather than providing an established mechanism.

While the present findings indicate an association between mind wandering and ADHD-related symptoms, they should be interpreted in light of how these constructs were measured. The MWQ primarily captures a general tendency toward everyday inattentiveness (e.g., *I have difficulty maintaining focus on simple or repetitive work.; I do things without paying full attention.)*, and several ASRS Inattention items describe closely related content (e.g., *How often do you have difficulty keeping your attention when you are doing boring or repetitive work?*; *How often do you make careless mistakes when you have to work on a boring or difficult project?)*. To evaluate this concern, we conducted supplementary analyses examining the association between ASRS and MWQ after separating Inattention from Hyperactivity/Impulsivity subscales, excluding the ASRS items with the closest semantic overlap to MWQ content, and modeling the overlapping and non-overlapping items simultaneously. These analyses indicated that item-content overlap contributes to, but does not fully account for, the observed association. A significant relationship with mind wandering was observed even for ASRS content not directly semantically related to MWQ wording, including the Hyperactivity/Impulsivity subscale and the non-overlapping ASRS items modeled both independently and alongside overlapping items.

The MWQ, as a static measure of general mind-wandering propensity, fails to capture other dimensions or content of mind wandering. Reducing it to a unidimensional construct may be problematic, as mind wandering has been increasingly recognized as a heterogeneous construct, characterized along dimensions such as task-relatedness, stimulus-dependence, intentionality, freedom of movement, valence, temporal orientation, goal-orientedness, and social aspects [67–71]. Focusing exclusively on frequency and not other dimensions has direct implications for the comparability of the present findings. Prior work has linked ADHD symptomatology specifically to spontaneous rather than deliberate mind wandering [72, 73]. As the present study assessed only general mind-wandering propensity, direct comparison with this literature is limited. Moreover, using task-embedded experience sampling Van den Driessche et al. [74] measured mind wandering separately from mind blanking - a state with no reportable mental content. They found that unmedicated children with ADHD and adults with subclinical ADHD showed no significant difference in mind wandering compared to controls, but rather increased mind blanking. This suggests that measuring mind wandering across multiple dimensions, rather than frequency alone, may reveal more complex associations with ADHD than the present cross-sectional unidimensional approach can capture.

More broadly, it remains unclear whether different psychopathological dimensions primarily differ in the overall propensity for mind wandering or in its phenomenological aspects, such as temporal orientation, emotional valence, and intentionality [21, 75, 76]. Based on our results, we might speculate that the propensity to disengage from the external environment, as captured by trait-level mind wandering measures such as the MWQ, may primarily reflect more general attentional control difficulties. However, the qualitative aspects of internally oriented attention, detectable with more dimensional scales, may vary across symptom dimensions, manifesting as ruminative, past-oriented thoughts in depression, worry-focused and future-oriented thoughts in anxiety, and intrusive, perseverative thoughts in OCD. From this perspective, ADHD-related attentional dysregulation may determine how frequently attention wanders from the task at hand, while other symptom dimensions shape the phenomenological nature of mind wandering episodes. This remains speculative, however, as the present study did not assess the content or phenomenology of mind wandering directly. Future work would benefit from more fine-grained methodological approaches, such as experience sampling, which could provide more detailed insight into the dynamics and content of mind wandering across different psychopathological dimensions and further distinguish mind wandering from ADHD-related inattentiveness at the construct level.

Our main regression analysis results also suggest that HCL-32, indicating hypomania symptoms, is a significant positive predictor of mind wandering. However, Bayesian regression provided no evidence favoring the alternative over the null hypothesis for this predictor. Results regarding HCL-32 also remained mixed across all supplementary analyses and the extended dataset, showing either non-significant or weak, inconsistent associations accounting for a negligible proportion of explained variance in mind wandering scores. Given this inconsistency, we abstain from overinterpreting this result. Existing literature offers no substantial empirical support or established theoretical accounts linking mind wandering and hypomania. One plausible contributing factor, however, is phenomenological overlap with “racing thoughts”. Racing thoughts are a hypomanic symptom describing a fast-moving, hard-to-control stream of thought. While racing thoughts and mind wandering have been shown to be empirically distinguishable constructs [77], their conceptual similarity with regard to the hardly controllable expression of spontaneous thoughts could plausibly introduce weak, inconsistent association observed here. Future studies could include measures that directly distinguish mind wandering from racing thoughts to clarify whether this overlap contributes to the present findings.

In sum, the present findings show that mind wandering was associated with a broad range of psychopathology-related symptom dimensions at the bivariate level, but only ADHD symptoms showed a consistently significant positive association once these dimensions were modeled jointly. This pattern proved robust across frequentist and Bayesian analyses, and in supplementary analyses addressing ASRS-MWQ item-content overlap and using extended samples including diagnosed and medicated participants. Whether our findings reflect a genuinely specific association with ADHD or a particularly strong marker of shared psychopathology-related variance remains an open question. Future work using latent-variable approaches and direct measures of attentional and executive functioning could help resolve it.

## Declarations

### Ethics approval and consent to participate

Participants provided informed consent, and the studies received approval from the Research Ethics Committee of Eötvös Loránd University, Budapest, Hungary (2024/214), in accordance with the principles of the Declaration of Helsinki.

### Data availability

The processed data and analysis code are available on the Open Science Framework (OSF) at https://doi.org/10.17605/OSF.IO/R48CQ.

### Author contributions

M.L.: Conceptualization, Methodology, Formal analysis, Data curation, Visualization, Writing – original draft. B.B.: Investigation, Data curation, Methodology, Writing – review & editing. T.V.: Conceptualization, Methodology, Software, Validation, Writing – review & editing. P.S.: Conceptualization, Supervision, Writing – review & editing. D.N.: Conceptualization, Supervision, Funding acquisition, Writing – review & editing.

## Acknowledgements

Not applicable.

