## Supplementary materials for "Disentangling the Relationship Between Mind Wandering and Symptom Dimensions in a Non-Clinical Sample: The Key Link with ADHD"

**S1. Sensitivity Analysis: ASRS Inattention and Hyperactivity/Impulsivity Subscales**

To examine whether the association between ADHD symptoms and mind wandering was specific to attentional symptom content, the ASRS Inattention and Hyperactivity/Impulsivity subscales were entered separately in place of the ASRS total score in the primary regression model.

**Table S1.**

*Regression Results for ASRS Inattention and Hyperactivity/Impulsivity Subscales Predicting Mind Wandering*

|  | **MWQ (mind wandering)** | | | | | | |
| --- | --- | --- | --- | --- | --- | --- | --- |
| *Predictors* | *β* | *SE* | *CI* | *t-value* | *p* | *df* | *sr²* |
| (Intercept) | 0.01 | 0.05 | -0.08 – 0.11 | 0.29 | 0.771 | 372.00 | - |
| ASRS (ADHD) - hyperactivity/impulsivity | 0.16 | 0.06 | 0.05 – 0.27 | 2.87 | **0.004** | 372.00 | .01 |
| ASRS (ADHD) - inattention | 0.43 | 0.06 | 0.32 – 0.54 | 7.51 | **<0.001** | 372.00 | .10 |
| MSS-B (schizotypy) | -0.10 | 0.06 | -0.22 – 0.02 | -1.70 | 0.090 | 372.00 | <.01 |
| ASQ (autism spectrum) | 0.03 | 0.05 | -0.07 – 0.13 | 0.56 | 0.574 | 372.00 | <.01 |
| OCI-R (OCD) | 0.06 | 0.05 | -0.04 – 0.17 | 1.15 | 0.249 | 372.00 | <.01 |
| BDI shortened (depression) | 0.01 | 0.06 | -0.11 – 0.13 | 0.16 | 0.873 | 372.00 | <.01 |
| HCL-32 (hypomania) | 0.09 | 0.05 | 0.00 – 0.18 | 2.03 | **0.043** | 372.00 | <.01 |
| EAT (eating attitude) | -0.01 | 0.05 | -0.11 – 0.08 | -0.30 | 0.766 | 372.00 | <.01 |
| Age | 0.02 | 0.04 | -0.07 – 0.10 | 0.38 | 0.706 | 372.00 | <.01 |
| Gender [male] | -0.09 | 0.11 | -0.31 – 0.12 | -0.83 | 0.406 | 372.00 | <.01 |
| Gender [other] | 0.96 | 0.59 | -0.21 – 2.13 | 1.62 | 0.106 | 372.00 | <.01 |
| Observations | 384 | | | | | | |
| R^2^ / R^2^ adjusted | 0.33 / 0.31 | | | | | | |
| *F_(11, 372)_ =* 16.29*, p <* .001 | | | | | | | |

*Note.* Standardized regression coefficients (β), standard errors (SE), 95% confidence intervals (CI), t-values, p-values, degrees of freedom, and semi-partial r² are reported for each predictor. p-values of significant predictors are indicated in bold. ASRS = Adult ADHD Self-Report Scale; MSS-B = Multidimensional Schizotypy Scale-Brief; ASQ = Autism Spectrum Quotient; OCI-R = Obsessive-Compulsive Inventory-Revised; BDI = Beck Depression Inventory; HCL-32 = Hypomania Checklist; EAT = Eating Attitude Test. Gender was dummy-coded with female as the reference category.

**S2. Sensitivity Analysis: ASRS-MWQ Item Content Overlap**

Given semantic overlap between ASRS and MWQ item content, we identified the ASRS items most closely related in content to the MWQ and examined their contribution to the ADHD-mind wandering association in two complementary analyses:

(a) excluding these items from the ASRS total score, and
(b) entering them alongside the remaining ASRS items simultaneously.

Identified items in ASRS score were the following:

- I7: How often do you make careless mistakes when you have to work on a boring or difficult project? (mean MWQ correlation coefficient: r = 0.32)

- I8: How often do you have difficulty keeping your attention when you are doing boring or repetitive work? (mean MWQ correlation coefficient: r = 0.44)

- I9: How often do you have difficulty concentrating on what people say to you, even when they are speaking to you directly? (mean MWQ correlation coefficient: r = 0.31)

**Table S2a.**

*Regression Results Excluding ASRS Items with Closest Semantic Overlap to MWQ Content*

|  | **MWQ (mind wandering)** | | | | | | |
| --- | --- | --- | --- | --- | --- | --- | --- |
| *Predictors* | *β* | *SE* | *CI* | *t-value* | *p* | *df* | *sr²* |
| (Intercept) | 0.03 | 0.05 | -0.07 – 0.12 | 0.50 | 0.619 | 373.00 | - |
| ASRS (ADHD) - low-overlap items | 0.42 | 0.05 | 0.32 – 0.53 | 7.73 | **<0.001** | 373.00 | .12 |
| MSS-B (schizotypy) | -0.07 | 0.06 | -0.20 – 0.05 | -1.22 | 0.224 | 373.00 | <.01 |
| ASQ (autism spectrum) | 0.05 | 0.05 | -0.06 – 0.16 | 0.90 | 0.367 | 373.00 | <.01 |
| OCI-R (OCD) | 0.06 | 0.06 | -0.05 – 0.17 | 1.07 | 0.284 | 373.00 | <.01 |
| BDI shortened (depression) | 0.06 | 0.06 | -0.06 – 0.18 | 1.03 | 0.306 | 373.00 | <.01 |
| HCL-32 (hypomania) | 0.12 | 0.05 | 0.03 – 0.21 | 2.57 | **0.010** | 373.00 | .01 |
| EAT (eating attitude) | -0.01 | 0.05 | -0.11 – 0.08 | -0.30 | 0.764 | 373.00 | <.01 |
| Age | 0.01 | 0.05 | -0.08 – 0.10 | 0.16 | 0.871 | 373.00 | <.01 |
| Gender [male] | -0.15 | 0.11 | -0.38 – 0.07 | -1.31 | 0.190 | 373.00 | .01 |
| Gender [other] | 1.18 | 0.62 | -0.04 – 2.40 | 1.90 | 0.058 | 373.00 | .01 |
| Observations | 384 | | | | | |  |
| R^2^ / R^2^ adjusted | 0.26 / 0.24 | | | | | |  |
| *F_(10, 373)_ =* 13.16*, p <* .001 | | | | | | | |

*Note.* Standardized regression coefficients (β), standard errors (SE), 95% confidence intervals (CI), t-values, p-values, degrees of freedom, and semi-partial r² are reported for each predictor. p-values of significant predictors are indicated in bold. ASRS = Adult ADHD Self-Report Scale; MSS-B = Multidimensional Schizotypy Scale-Brief; ASQ = Autism Spectrum Quotient; OCI-R = Obsessive-Compulsive Inventory-Revised; BDI = Beck Depression Inventory; HCL-32 = Hypomania Checklist; EAT = Eating Attitude Test. Gender was dummy-coded with female as the reference category.

**Table S2b.**

*Regression Results with ASRS Overlapping and Remaining Items Entered Simultaneously*

|  | **MWQ (mind wandering)** | | | | | | |
| --- | --- | --- | --- | --- | --- | --- | --- |
| *Predictors* | *β* | *SE* | *CI* | *t-value* | *p* | *df* | *sr²* |
| (Intercept) | 0.01 | 0.05 | -0.08 – 0.10 | 0.18 | 0.859 | 372.00 | - |
| ASRS (ADHD) - high-overlap items | 0.46 | 0.05 | 0.35 – 0.57 | 8.40 | **<0.001** | 372.00 | .12 |
| ASRS (ADHD) - low-overlap items | 0.20 | 0.06 | 0.09 – 0.31 | 3.52 | **<0.001** | 372.00 | .02 |
| MSS-B (schizotypy) | -0.10 | 0.06 | -0.21 – 0.01 | -1.73 | 0.084 | 372.00 | <.01 |
| ASQ (autism spectrum) | 0.01 | 0.05 | -0.09 – 0.11 | 0.22 | 0.825 | 372.00 | <.01 |
| OCI-R (OCD) | 0.05 | 0.05 | -0.05 – 0.16 | 1.04 | 0.298 | 372.00 | <.01 |
| BDI shortened (depression) | -0.01 | 0.06 | -0.12 – 0.10 | -0.15 | 0.880 | 372.00 | <.01 |
| HCL-32 (hypomania) | 0.07 | 0.04 | -0.02 – 0.15 | 1.53 | 0.128 | 372.00 | <.01 |
| EAT (eating attitude) | -0.01 | 0.04 | -0.10 – 0.07 | -0.30 | 0.764 | 372.00 | <.01 |
| Age | 0.01 | 0.04 | -0.07 – 0.09 | 0.24 | 0.810 | 372.00 | <.01 |
| Gender [male] | -0.06 | 0.11 | -0.27 – 0.15 | -0.56 | 0.578 | 372.00 | <.01 |
| Gender [other] | 0.77 | 0.57 | -0.35 – 1.89 | 1.35 | 0.178 | 372.00 | <.01 |
| Observations | 384 | | | | | | |
| R^2^ / R^2^ adjusted | 0.38 / 0.36 | | | | | | |
| *F_(11, 372)_ =* 20.61*, p <* .001 | | | | | | | |

*Note.* Standardized regression coefficients (β), standard errors (SE), 95% confidence intervals (CI), t-values, p-values, degrees of freedom, and semi-partial r² are reported for each predictor. p-values of significant predictors are indicated in bold. ASRS = Adult ADHD Self-Report Scale; MSS-B = Multidimensional Schizotypy Scale-Brief; ASQ = Autism Spectrum Quotient; OCI-R = Obsessive-Compulsive Inventory-Revised; BDI = Beck Depression Inventory; HCL-32 = Hypomania Checklist; EAT = Eating Attitude Test. Gender was dummy-coded with female as the reference category.

**S3. Sensitivity Analysis: ASRS × Data Collection Round Interaction**

Given that the MWQ response format differed across the two data collection rounds, we tested an ASRS × Round interaction in the primary regression model to assess whether the association between ADHD symptoms and mind wandering was consistent across rounds.

**Table S3.**

*Regression Results for the ASRS × Data Collection Round Interaction*

|  | **MWQ (mind wandering)** | | | | | |
| --- | --- | --- | --- | --- | --- | --- |
| *Predictors* | *β* | *SE* | *CI* | *t-value* | *p* | *df* |
| (Intercept) | 0.00 | 0.08 | -0.15 – 0.15 | 0.02 | 0.984 | 371.00 |
| ASRS (ADHD) | 0.46 | 0.08 | 0.30 – 0.61 | 5.73 | **<0.001** | 371.00 |
| MSS-B (schizotypy) | -0.11 | 0.06 | -0.22 – 0.01 | -1.76 | 0.079 | 371.00 |
| ASQ (autism spectrum) | 0.04 | 0.05 | -0.06 – 0.14 | 0.73 | 0.463 | 371.00 |
| OCI-R (OCD) | 0.03 | 0.06 | -0.09 – 0.14 | 0.48 | 0.635 | 371.00 |
| BDI shortened (depression) | 0.04 | 0.06 | -0.08 – 0.16 | 0.68 | 0.500 | 371.00 |
| HCL-32 (hypomania) | 0.10 | 0.05 | 0.01 – 0.19 | 2.11 | **0.036** | 371.00 |
| EAT (eating attitude) | -0.01 | 0.05 | -0.10 – 0.08 | -0.22 | 0.827 | 371.00 |
| Age | 0.01 | 0.04 | -0.08 – 0.09 | 0.13 | 0.893 | 371.00 |
| Gender [male] | -0.12 | 0.11 | -0.34 – 0.10 | -1.09 | 0.275 | 371.00 |
| Gender [other] | 1.02 | 0.60 | -0.16 – 2.20 | 1.70 | 0.091 | 371.00 |
| Data round | 0.03 | 0.10 | -0.16 – 0.22 | 0.33 | 0.739 | 371.00 |
| ASRS (ADHD) × Data round | 0.10 | 0.09 | -0.08 – 0.28 | 1.10 | 0.271 | 371.00 |
| Observations | 384 | | | | | |
| R^2^ / R^2^ adjusted | 0.31 / 0.29 | | | | | |
| *F_(12, 371)_ =* 14.12*, p <* .001 | | | | | | |

*Note.* Standardized regression coefficients (β), standard errors (SE), 95% confidence intervals (CI), t-values, p-values, and degrees of freedom are reported for each predictor. p-values of significant predictors are indicated in bold. ASRS = Adult ADHD Self-Report Scale; MSS-B = Multidimensional Schizotypy Scale-Brief; ASQ = Autism Spectrum Quotient; OCI-R = Obsessive-Compulsive Inventory-Revised; BDI = Beck Depression Inventory; HCL-32 = Hypomania Checklist; EAT = Eating Attitude Test. Gender was dummy-coded with female as the reference category.

Round-specific ADHD-mind wandering slopes were highly comparable: Round 1: β = 0.46, 95% CI [0.30, 0.61]; Round 2: β = 0.56, 95% CI [0.43, 0.69]. The direct contrast between rounds was non-significant (difference = −0.101, *SE* = 0.09, *t(371)* = −1.10, *p* = .271), confirming the ASRS × Round interaction result. In summary, the association between ADHD symptoms and mind wandering did not differ meaningfully between data collection rounds.

**S4. Sensitivity Analysis: Prior Specification Robustness**

To assess the robustness of our findings to prior specification, we conducted sensitivity analyses using two alternative prior distributions for the regression coefficients in the Bayesian multiple linear regression model predicting mind wandering from psychopathology-related measures.

The main analysis (reported in the manuscript) used weakly informative priors [Normal(0, 1)] for regression coefficients, which provide slight regularization without strongly constraining the estimates. Here we report results using:

- Moderately conservative priors for regression coefficients - Normal(0, 0.5)
- Strongly regularizing priors for regression coefficients: Normal(0, 0.3)

These alternative priors progressively increase regularization, shrinking coefficient estimates toward zero. Intercept prior [Normal(0, 1)] was identical across all specifications. Consistency of findings across these specifications would indicate that our conclusions are not unduly influenced by prior choice.

**Table S4**

*Bayesian multiple linear regression with moderately conservative priors for regression coefficients [Normal(0, 0.5)]*

|  | **MWQ (mind wandering)** | | | |
| --- | --- | --- | --- | --- |
| **Predictor** | **Estimate** | **Est. Error** | **95% CrI** | **BF₁₀** |
| ASRS (ADHD) | 0.52 | 0.05 | [0.41, 0.62] | > 1000 |
| MSS-B (schizotypy) | -0.10 | 0.06 | [-0.21, 0.02] | 0.44 |
| ASQ (autism spectrum) | 0.04 | 0.05 | [-0.07, 0.14] | 0.13 |
| OCI-R (OCD) | 0.04 | 0.05 | [-0.07, 0.15] | 0.14 |
| BDI shortened (depression) | 0.03 | 0.06 | [-0.08, 0.15] | 0.14 |
| HCL-32 (hypomania) | 0.09 | 0.05 | [0.01, 0.18] | 0.75 |
| EAT (eating attitude) | -0.01 | 0.05 | [-0.11, 0.08] | 0.10 |
| Age | 0.00 | 0.04 | [-0.08, 0.09] | 0.09 |
| Gender [male] | -0.12 | 0.11 | [-0.34, 0.09] | 0.40 |
| Gender [other] | 0.42 | 0.38 | [-0.34, 1.17] | 1.40 |
| Bayesian R² = .31, 95% CrI [0.24, 0.37] | | | | |

*Note.* Standardized posterior means (β), standard errors of the posterior distribution (Est.Error), 95% credible intervals (CrI), and Bayes factors (BF₁₀) are reported for each predictor. BF₁₀ values indicate evidence strength: <1 (evidence for null), 1-3 (anecdotal evidence for alternative), 3-10 (moderate evidence for alternative), 10-100 (strong evidence for alternative), >100 (decisive evidence for alternative). ASRS = Adult ADHD Self-Report Scale; MSS-B = Multidimensional Schizotypy Scale-Brief; ASQ = Autism Spectrum Quotient; OCI-R = Obsessive-Compulsive Inventory-Revised; BDI = Beck Depression Inventory; HCL-32 = Hypomania Checklist; EAT = Eating Attitude Test. Gender was dummy-coded with female as the reference category.

**Table S5**

*Bayesian multiple linear regression with strongly regularizing priors for regression coefficients [Normal(0, 0.3)]*

|  | **MWQ (mind wandering)** | | | |
| --- | --- | --- | --- | --- |
| **Predictor** | **Estimate** | **Est. Error** | **95% CrI** | **BF₁₀** |
| ASRS (ADHD) | 0.51 | 0.05 | [0.40, 0.61] | > 1000 |
| MSS-B (schizotypy) | -0.09 | 0.06 | [-0.21, 0.02] | 0.72 |
| ASQ (autism spectrum) | 0.03 | 0.05 | [-0.07, 0.13] | 0.21 |
| OCI-R (OCD) | 0.04 | 0.05 | [-0.06, 0.15] | 0.24 |
| BDI shortened (depression) | 0.04 | 0.06 | [-0.08, 0.15] | 0.23 |
| HCL-32 (hypomania) | 0.09 | 0.04 | [0.01, 0.18] | 1.34 |
| EAT (eating attitude) | -0.01 | 0.05 | [-0.10, 0.08] | 0.16 |
| Age | 0.00 | 0.04 | [-0.08, 0.09] | 0.14 |
| Gender [male] | -0.11 | 0.10 | [-0.32, 0.09] | 0.63 |
| Gender [other] | 0.21 | 0.27 | [-0.32, 0.73] | 1.21 |
| Bayesian R² = 0.30, 95% CrI [0.24, 0.36] | | | | |

*Note.* Standardized posterior means (β), standard errors of the posterior distribution (Est.Error), 95% credible intervals (CrI), and Bayes factors (BF₁₀) are reported for each predictor. BF₁₀ values indicate evidence strength: <1 (evidence for null), 1-3 (anecdotal evidence for alternative), 3-10 (moderate evidence for alternative), 10-100 (strong evidence for alternative), >100 (decisive evidence for alternative). ASRS = Adult ADHD Self-Report Scale; MSS-B = Multidimensional Schizotypy Scale-Brief; ASQ = Autism Spectrum Quotient; OCI-R = Obsessive-Compulsive Inventory-Revised; BDI = Beck Depression Inventory; HCL-32 = Hypomania Checklist; EAT = Eating Attitude Test. Gender was dummy-coded with female as the reference category.

### **S5. Extended Dataset Analysis**

The main analysis (reported in the manuscript) was conducted on participants without self-reported mental health diagnoses or psychotropic medication use (N = 384). We additionally conducted a re-analysis using the extended dataset that included all participants (N = 441). This extended dataset includes the 57 participants who were excluded from the main analysis due to a self-reported mental health diagnosis and/or active use of psychotropic medication. Characteristics regarding the reported diagnoses and medication status of these 57 participants are reported below in Table S6.

**Table S6**

*Characteristics of participants excluded from the primary sample based on self-reported conditions and/or psychotropic medication use (N = 57)*

| **Category** | **Self-reported condition / medication status** | **n (%)** |
| --- | --- | --- |
| **A. Self-reported conditions** |  |  |
| Neurological history | Neurological condition or Epilepsy | 4 (7) |
| Neurodevelopmental conditions | Autism spectrum disorder (ASD) | 2 (3.5) |
|  | Attention-deficit/hyperactivity disorder (ADHD) | 4 (7) |
| Mental health conditions | Obsessive-compulsive disorder (OCD) | 1 (1.7) |
|  | Schizophrenia | 1 (1.7) |
|  | Depression | 19 (33.3) |
|  | Anxiety disorder | 22 (38.6) |
|  | Eating disorder (ED) | 4 (7) |
|  | Personality disorder (PD) | 1 (1.7) |
| **B. Psychotropic and recreational medication use** |  |  |
|  | Recreational | 1 (1.7) |
|  | Psychotropic | 13 (22.8) |

*Note.* Participants could report more than one condition and/or medication

**S5.1. Descriptive statistics**

**Table S7**

*Descriptive statistics and internal consistency for all study variables*

| **Questionnaire** | **Mean (SD)** | **Skewness** | **Kurtosis** | **Cronbach's α** |
| --- | --- | --- | --- | --- |
| MWQ (mind wandering) (Round 1) | 19.19 (7.44) | −0.38 | −0.17 | 0.88 |
| MWQ (mind wandering) (Round 2) | 20.05 (5.17) | −0.16 | −0.62 | 0.82 |
| ASQ (autism spectrum) | 19.24 (5.84) | 0.38 | −0.06 | 0.80 |
| ASRS (ADHD) | 50.55 (9.52) | 0.12 | 0.02 | 0.84 |
| BDI shortened (depression) | 14.87 (5.08) | 0.71 | −0.39 | 0.87 |
| EAT (eating attitude) | 12.02 (11.58) | 1.56 | 2.32 | 0.90 |
| HCL-32 (hypomania) | 18.08 (6.20) | −0.60 | 0.46 | 0.87 |
| MSS-B (schizotypy) | 8.66 (6.79) | 0.93 | 0.25 | 0.89 |
| OCI-R (OCD) | 20.42 (13.08) | 0.40 | −0.65 | 0.91 |
| **Age:** *M =* 22.2, *SD =* 4.1, range = 18 - 52  **Gender:** F = 347, M = 91, Other = 3 | | | | |

*Note.* Mean scores, standard deviations, skewness, kurtosis, and Cronbach's alpha coefficients are presented for each questionnaire in the extended dataset (N = 441). MWQ statistics are reported separately by data collection round due to different response scales used (Round 1: N = 168; Round 2: N = 273). ASQ = Autism Spectrum Quotient; ASRS = Adult ADHD Self-Report Scale; BDI = Beck Depression Inventory; EAT = Eating Attitude Test; HCL-32 = Hypomania Checklist; MSS-B = Multidimensional Schizotypy Scale-Brief; MWQ = Mind-Wandering Questionnaire; OCI-R = Obsessive-Compulsive Inventory-Revised.

**S5.2. Bivariate correlation analysis**

**Figure S1**

*Correlations among mind wandering and psychopathology-related measures*

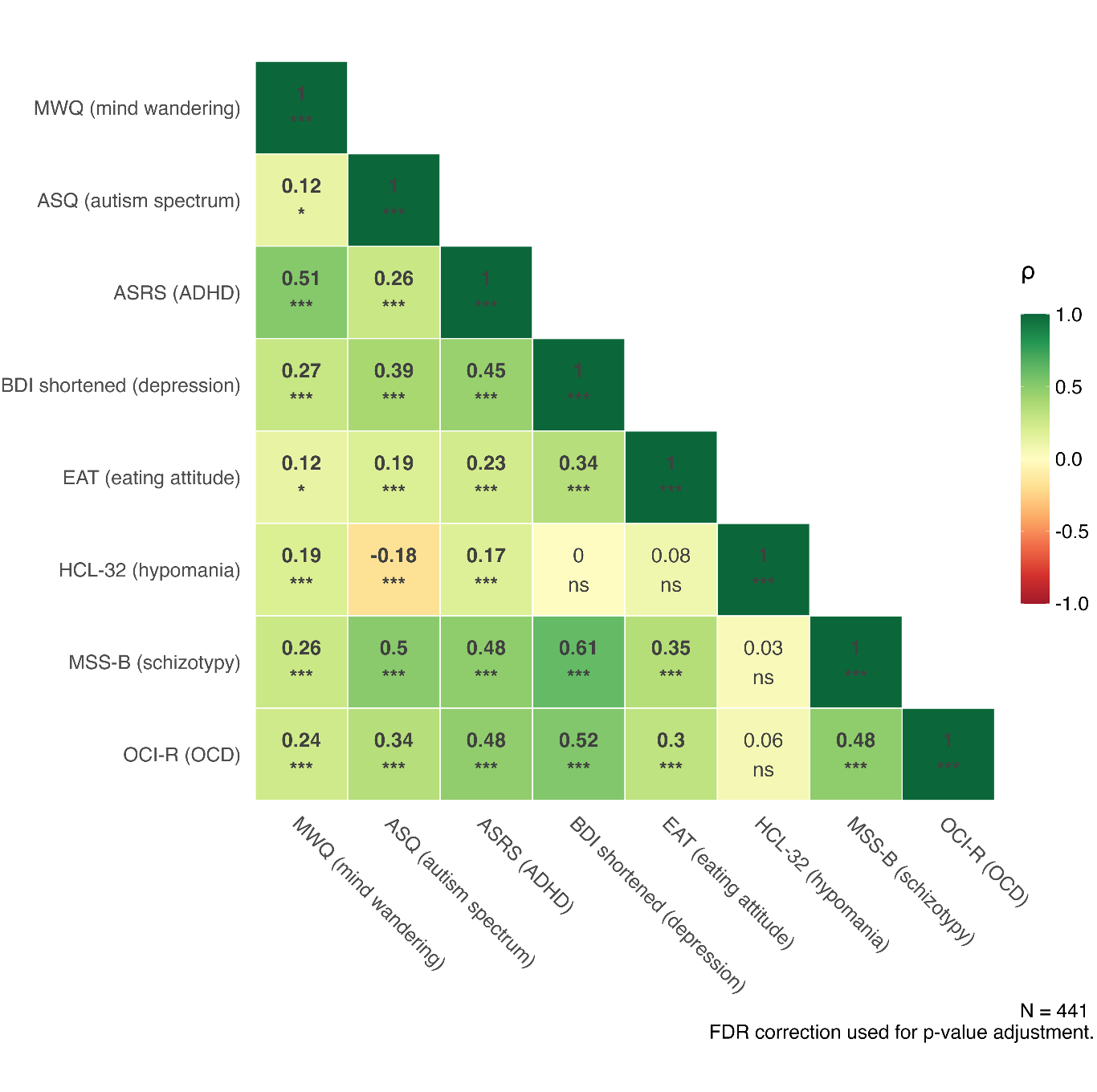
*Note.* ASQ = Autism Spectrum Quotient; ASRS = Adult ADHD Self-Report Scale; BDI = Beck Depression Inventory; EAT = Eating Attitude Test; HCL-32 = Hypomania Checklist; MSS-B = Multidimensional Schizotypy Scale-Brief; MWQ = Mind-Wandering Questionnaire; OCI-R = Obsessive-Compulsive Inventory-Revised.

* p < .05; ** p < .01; *** p < .001 (FDR-corrected)

**S5.3. Multiple regression analysis**

**Table S8**

*Multiple linear regression predicting mind wandering from psychopathology-related measures*

|  | **MWQ (mind wandering)** | | | | | | |
| --- | --- | --- | --- | --- | --- | --- | --- |
| *Predictors* | *β* | *SE* | *CI* | *t-value* | *p* | *df* | *sr²* |
| (Intercept) | 0.02 | 0.05 | -0.07 – 0.11 | 0.36 | 0.719 | 430.00 | - |
| ASRS (ADHD) | 0.51 | 0.05 | 0.41 – 0.60 | 10.13 | **<0.001** | 430.00 | .17 |
| MSS-B (schizotypy) | -0.11 | 0.06 | -0.22 – 0.00 | -1.93 | 0.055 | 430.00 | <.01 |
| ASQ (autism spectrum) | 0.04 | 0.05 | -0.06 – 0.13 | 0.74 | 0.461 | 430.00 | <.01 |
| OCI-R (OCD) | -0.00 | 0.05 | -0.10 – 0.10 | -0.04 | 0.969 | 430.00 | <.01 |
| BDI shortened (depression) | 0.09 | 0.06 | -0.02 – 0.20 | 1.61 | 0.108 | 430.00 | <.01 |
| HCL-32 (hypomania) | 0.11 | 0.04 | 0.02 – 0.19 | 2.47 | **0.014** | 430.00 | <.01 |
| EAT (eating attitude) | -0.01 | 0.04 | -0.10 – 0.07 | -0.30 | 0.767 | 430.00 | <.01 |
| Age | -0.00 | 0.04 | -0.08 – 0.08 | -0.06 | 0.949 | 430.00 | <.01 |
| Gender [male] | -0.09 | 0.10 | -0.29 – 0.12 | -0.83 | 0.405 | 430.00 | <.01 |
| Gender [other] | 0.20 | 0.50 | -0.77 – 1.18 | 0.41 | 0.682 | 430.00 | <.01 |
| Observations | 441 | | | | | | |
| R^2^ / R^2^ adjusted | .30 / .28 | | | | | | |
| F_(10, 430)_ = 18.05, p < .001 | | | | | | | |

*Note.* Standardized regression coefficients (β), standard errors (SE), 95% confidence intervals (CI), t-values, p-values, degrees of freedom, and semi-partial r² are reported for each predictor. p-values of significant predictors are indicated in bold. ASRS = Adult ADHD Self-Report Scale; MSS-B = Multidimensional Schizotypy Scale-Brief; ASQ = Autism Spectrum Quotient; OCI-R = Obsessive-Compulsive Inventory-Revised; BDI = Beck Depression Inventory; HCL-32 = Hypomania Checklist; EAT = Eating Attitude Test. Gender was dummy-coded with female as the reference category.

**Figure S2**

*Multiple linear regression predicting mind wandering from psychopathology-related measures*

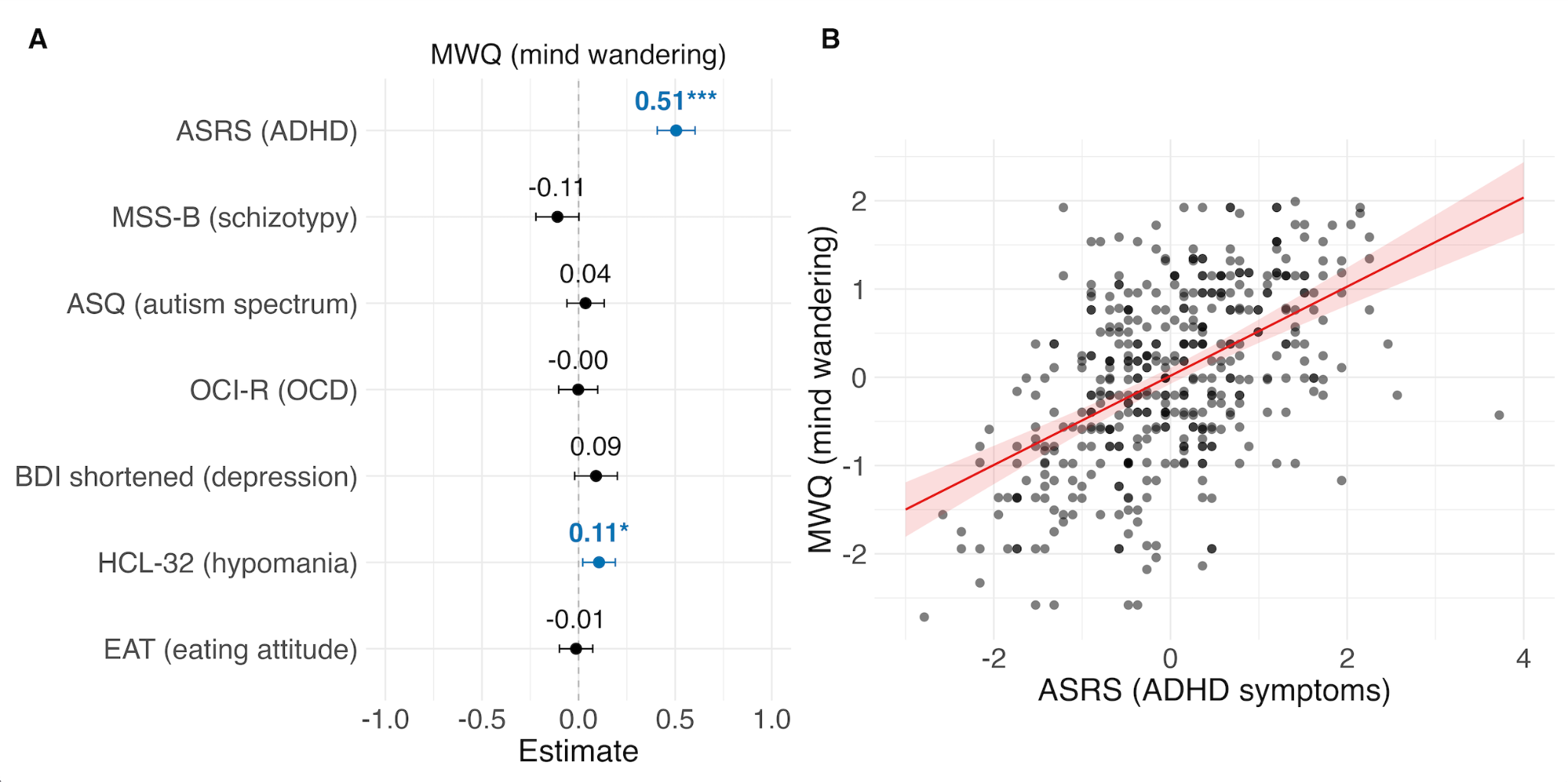

*Note.* **Panel A:** Standardized regression coefficients (β) with 95% confidence intervals for all predictors. The vertical dashed line represents no effect (β = 0). **Panel B:** Model-based predicted mind wandering scores as a function of ADHD (ASRS), estimated from a multiple regression model. Predictions are shown with all other predictors held constant at their mean. Individual data points represent standardized scores, the red line shows the predicted linear relationship, and the shaded ribbon represents the 95% confidence interval around the prediction line. ASRS = Adult ADHD Self-Report Scale; MSS-B = Multidimensional Schizotypy Scale-Brief; ASQ = Autism Spectrum Quotient; OCI-R = Obsessive-Compulsive Inventory-Revised; BDI = Beck Depression Inventory; HCL-32 = Hypomania Checklist; EAT = Eating Attitude Test; MWQ = Mind-Wandering Questionnaire.

**S5.4. Bayesian Multiple Linear Regression**

We conducted Bayesian multiple linear regression analyses on the extended dataset using three prior specifications: weakly informative [Normal(0, 1)], moderately conservative [Normal(0, 0.5)], and strongly regularizing [Normal(0, 0.3)] priors for regression coefficients. Intercept prior [Normal(0, 1)] was identical across all specifications.

**Table S9**

*Bayesian multiple linear regression with weakly informative priors for regression coefficients [Normal(0, 1)]*

|  | **MWQ (mind wandering)** | | | |
| --- | --- | --- | --- | --- |
| **Predictor** | **Estimate** | **Est. Error** | **95% CrI** | **BF₁₀** |
| ASRS (ADHD) | 0.50 | 0.05 | [0.41, 0.60] | > 1000 |
| MSS-B (schizotypy) | -0.11 | 0.06 | [-0.22, 0.00] | 0.36 |
| ASQ (autism spectrum) | 0.04 | 0.05 | [-0.06, 0.13] | 0.06 |
| OCI-R (OCD) | 0.00 | 0.05 | [-0.10, 0.10] | 0.05 |
| BDI shortened (depression) | 0.09 | 0.06 | [-0.02, 0.20] | 0.21 |
| HCL-32 (hypomania) | 0.11 | 0.04 | [0.02, 0.19] | 0.93 |
| EAT (eating attitude) | -0.01 | 0.04 | [-0.10, 0.07] | 0.05 |
| Age | 0.00 | 0.04 | [-0.08, 0.08] | 0.04 |
| Gender [male] | -0.09 | 0.10 | [-0.29, 0.12] | 0.14 |
| Gender [other] | 0.16 | 0.45 | [-0.71, 1.04] | 0.48 |
| Bayesian R² = .30, 95% CrI [0.24, 0.36] | | | | |

*Note.* Standardized posterior means (β), standard errors of the posterior distribution (Est.Error), 95% credible intervals (CrI), and Bayes factors (BF₁₀) are reported for each predictor. BF₁₀ values indicate evidence strength: <1 (evidence for null), 1-3 (anecdotal evidence for alternative), 3-10 (moderate evidence for alternative), 10-100 (strong evidence for alternative), >100 (decisive evidence for alternative). ASRS = Adult ADHD Self-Report Scale; MSS-B = Multidimensional Schizotypy Scale-Brief; ASQ = Autism Spectrum Quotient; OCI-R = Obsessive-Compulsive Inventory-Revised; BDI = Beck Depression Inventory; HCL-32 = Hypomania Checklist; EAT = Eating Attitude Test. Gender was dummy-coded with female as the reference category.

**Table S10**

*Bayesian multiple linear regression with moderately conservative priors for regression coefficients [Normal(0, 0.5)]*

|  | **MWQ (mind wandering)** | | | |
| --- | --- | --- | --- | --- |
| **Predictor** | **Estimate** | **Est. Error** | **95% CrI** | **BF₁₀** |
| ASRS (ADHD) | 0.50 | 0.05 | [0.40, 0.60] | > 1000 |
| MSS-B (schizotypy) | -0.11 | 0.06 | [-0.22, 0.00] | 0.67 |
| ASQ (autism spectrum) | 0.03 | 0.05 | [-0.06, 0.13] | 0.13 |
| OCI-R (OCD) | 0.00 | 0.05 | [-0.10, 0.10] | 0.10 |
| BDI shortened (depression) | 0.09 | 0.06 | [-0.02, 0.20] | 0.40 |
| HCL-32 (hypomania) | 0.11 | 0.04 | [0.02, 0.19] | 1.89 |
| EAT (eating attitude) | -0.01 | 0.04 | [-0.10, 0.07] | 0.09 |
| Age | 0.00 | 0.04 | [-0.08, 0.08] | 0.08 |
| Gender [male] | -0.08 | 0.10 | [-0.28, 0.12] | 0.29 |
| Gender [other] | 0.10 | 0.35 | [-0.59, 0.79] | 0.73 |
| Bayesian R² = .30, 95% CrI [0.24, 0.35] | | | | |

*Note.* Standardized posterior means (β), standard errors of the posterior distribution (Est.Error), 95% credible intervals (CrI), and Bayes factors (BF₁₀) are reported for each predictor. BF₁₀ values indicate evidence strength: <1 (evidence for null), 1-3 (anecdotal evidence for alternative), 3-10 (moderate evidence for alternative), 10-100 (strong evidence for alternative), >100 (decisive evidence for alternative). ASRS = Adult ADHD Self-Report Scale; MSS-B = Multidimensional Schizotypy Scale-Brief; ASQ = Autism Spectrum Quotient; OCI-R = Obsessive-Compulsive Inventory-Revised; BDI = Beck Depression Inventory; HCL-32 = Hypomania Checklist; EAT = Eating Attitude Test. Gender was dummy-coded with female as the reference category.

**Table S11**

*Bayesian multiple linear regression with strongly regularizing priors for regression coefficients [Normal(0, 0.3)]*

|  | **MWQ (mind wandering)** | | | |
| --- | --- | --- | --- | --- |
| **Predictor** | **Estimate** | **Est. Error** | **95% CrI** | **BF₁₀** |
| ASRS (ADHD) | 0.49 | 0.05 | [0.40, 0.59] | > 1000 |
| MSS-B (schizotypy) | -0.10 | 0.05 | [-0.21, 0.01] | 1.02 |
| ASQ (autism spectrum) | 0.03 | 0.06 | [-0.06, 0.13] | 0.20 |
| OCI-R (OCD) | 0.00 | 0.05 | [-0.10, 0.10] | 0.17 |
| BDI shortened (depression) | 0.09 | 0.05 | [-0.02, 0.20] | 0.68 |
| HCL-32 (hypomania) | 0.11 | 0.06 | [0.02, 0.19] | 3.10 |
| EAT (eating attitude) | -0.01 | 0.04 | [-0.10, 0.07] | 0.15 |
| Age | 0.00 | 0.04 | [-0.08, 0.08] | 0.14 |
| Gender [male] | -0.08 | 0.04 | [-0.27, 0.11] | 0.45 |
| Gender [other] | 0.06 | 0.10 | [-0.45, 0.56] | 0.87 |
| Bayesian R² = .29, 95% CrI [0.23, 0.35] | | | | |

*Note.* Standardized posterior means (β), standard errors of the posterior distribution (Est.Error), 95% credible intervals (CrI), and Bayes factors (BF₁₀) are reported for each predictor. BF₁₀ values indicate evidence strength: <1 (evidence for null), 1-3 (anecdotal evidence for alternative), 3-10 (moderate evidence for alternative), 10-100 (strong evidence for alternative), >100 (decisive evidence for alternative). ASRS = Adult ADHD Self-Report Scale; MSS-B = Multidimensional Schizotypy Scale-Brief; ASQ = Autism Spectrum Quotient; OCI-R = Obsessive-Compulsive Inventory-Revised; BDI = Beck Depression Inventory; HCL-32 = Hypomania Checklist; EAT = Eating Attitude Test. Gender was dummy-coded with female as the reference category.
